# Compromised futures: gastrointestinal parasites reduce survival and recruitment but not reproductive performance in wild female spotted hyenas (*Crocuta crocuta*)

**DOI:** 10.64898/2025.12.16.694767

**Authors:** Susana Patrícia Veloso Soares, Miguel Mendes Veiga, Stephan Karl, Sonja Metzger, Gábor Árpád Czirják, Marion Linda East, Heribert Hofer, Jella Wauters, Susana Carolina Martins Ferreira, Morgane Gicquel, Sarah Benhaiem

## Abstract

**Background:** Gastrointestinal parasitic (GIP) infections can decrease the survival and reproductive performance of humans and livestock. Even so, the immediate and delayed fitness consequences of GIP in unmanaged wild mammal populations, particularly during the sensitive and energetically demanding early life stage, remain largely unknown.

**Methods:** We investigate non-invasively the effects of GIP during early life on the survival to adulthood, age at first reproduction and longevity of 68 individually known female spotted hyenas in the Serengeti National Park, Tanzania. We also test whether a high investment in gastrointestinal local immunity (i.e. faecal mucin and immunoglobulin A) and an elevated allostatic load (i.e. faecal glucocorticoid metabolites) decrease these measures of performance, when controlling for resource availability and maternal effects (i.e. maternal social rank) using logistic regression and Cox proportional hazards models.

**Results:** *Ancylostoma* sp. infections and the level of immunoglobulin A both negatively associate with survival to adulthood, probably because of the damage caused by the parasite and resource allocation trade-off between immunity and maintenance. Conversely, maternal social rank has a positive influence on survival to adulthood and longevity. Immune responses, allostatic load and GIP do not affect the age at first reproduction, suggesting either no effect, a complex interaction between different life stages, or a more immediate time-dependent effect.

**Conclusion:** Our study highlights the critical role of GIP infections and maternal effects on early-life fitness and long-term outcomes in females of a wild carnivore, thereby contributing to our understanding of ecological dynamics in unmanaged ecosystems.

## Background

Early life - a period often defined from conception to maturity [1] - is a critical period for mammalian development [2]. In wild mammals, experiences encountered during early life driven by environmental or climatic conditions [3,4] or parental effects such as social rank in group-living species [5,6] can have long-lasting effects on physiology, behaviour, morphology and ultimately health, performance and fitness [7–9]. In humans and livestock, sub-lethal gastrointestinal parasitic infections (GIP), i.e. with protozoans and helminths, can cause undernourishment and developmental impairments, especially when experienced during early life, with a particularly pronounced impact on females and offspring (e.g., humans [10,11]; livestock [12,13]). These compromises in the fulfillment of their reproductive/productive potential undermine subsequent generations from their very beginnings, perpetuating a vicious cycle of poor health, performance, and fitness. Recent field studies have linked GIP infections during early life to immediate performance measures such as overwinter survival in red deer, *Cervus elaphus* [14] or predation risk in leverets of Iberian hares, *Lepus granatensis* [15]. However, most free-ranging wildlife studies still rely on the artificial reduction of parasites [16,17] and are not focusing on the early life period [18,19]. There is also a general lack of studies evaluating both the short and long-term fitness costs of GIP infections during early life in unmanaged wildlife populations [20] nor is it clear whether these effects mirror those seen in humans and livestock.

Early life immunity can also render young individuals more susceptible to infections [21,22]. As a result, young mammals often harbour higher loads of parasites than adults [23,24]. While their developing immune systems are learning to efficiently recognize and regulate pathogens, front-line defenses of the gut, such as mucin and immunoglobulin A (IgA), play crucial roles in host-parasite interactions [25,26]. Mucin is an integrated component of the mucus layer that covers the intestinal epithelium and provides both physical and chemical protection against pathogens, modulating gut colonization [27,28]. IgA, the most abundant antibody in the gut, aids in pathogen removal, microbiome regulation, and inflammation control [29,30]. Producing these immune responses increases resource costs for the host which may reduce fitness [31,32]. However, the significance of these costs as a potential component of adversity during early life on fitness is rarely determined in unmanaged wildlife populations.

Another physiological aspect that may influence the immediate and delayed effects of/in early life is allostatic load. Allostasis is the process of maintaining physiological stability through behavioural or physiological responses to challenges, and its cumulative energetic cost is termed allostatic load [33,34]. The adjustment of mediators, such as glucocorticoids, enables the modulation of resources and energy, which is fundamental to addressing these challenges [35]. These mediators are involved in energy metabolism, provide significant protection during stages of elevated energetic demand, and contribute substantially to post-exposure recuperation [36]. The repeated (or multiple) exposure to challenges can lead to chronically elevated levels of these mediators that result in dysfunction of multiple somatic systems, causing context-dependent maladaptive changes such as appetite loss and immune/reproductive modulation [37]. Such dysfunction can impair individual fitness by reducing body condition, probability of survival, or reproductive output [38–40]. Because allostatic systems are still developing in young mammals, they are more malleable and vulnerable [41,42] and thus early life experiences may result in profound and lasting effects. However, the ontogenetic role and fitness costs of allostatic load in unmanaged wildlife GIP infections also remains understudied, as most research focuses on mature individuals (e.g. [43]).

Free-ranging spotted hyenas (*Crocuta crocuta*), hereafter hyenas, are a useful model for studying the short and long-term consequences of early-life GIP infections. The slow pace of their early life period, marked by extended gestation (110 days - reviewed by [44]), a protracted lactation period of 14-20 months, adulthood being typically set at 2 years [45,46] alongside a long lifespan in some individuals [47,48], allows for an observable gradual development and subsequent manifestation of the effects of GIP infections over time. Previous studies have shown that factors like maternal social rank or rainfall can influence fitness related measures in different populations over different life stages [48–51]. As a social apex carnivore, hyenas host diverse GIP species modulated by host age and some of the same ecological, demographic, physiological, and behavioural factors influencing fitness [24,52–54]. Past studies have also shown that resource allocation to immune responses (IgA) and high faecal egg loads of hookworms (*Ancylostoma sp*.) negatively affect juvenile survival to adulthood and longevity in spotted hyenas [24,26]. However, we have yet to understand how GIP, immunity, and allostasis during early life shape both short and long-term performance measures and their relative importance on each measure.

Here, we aim to investigate the effects of early-life GIP on short and long-term measures of performance in female hyenas in the Serengeti studied in the context of a long-term project. As part of this project, faecal samples from individually known animals are opportunistically collected and analysed routinely for proxies of the presence and abundance of GIP, allostatic load and two markers of the local mucosal immune response (mucin and IgA) [24,26,55]. We focus on the transition to adulthood and the start of the reproductive career, considering survival to adulthood and age of first reproduction as measures of performance. We also look into a longer perspective, considering longevity. We examine the effects of *Ancylostoma* sp.eggs loads, parasitic co-infections - hereafter polyparasitism, faecal mucin, faecal-IgA and allostatic load (faecal glucocorticoid metabolites), while controlling for a key environmental influence - maternal social rank. Maternal social rank is an important factor in this species: it determines a mother’s access to resources and as a result offspring growth rate, survival to adulthood and age at first reproduction [56,48]. Because early life is a highly sensitive, demanding, and complex life stage, we expect GIP to exert both an immediate and delayed negative effect on their hosts. Activation of the immune system and chronically elevated allostatic load may further increase energetic costs for organisms, leading to early life-history trade-offs. These trade-offs may negatively affect short-term fitness-linked performance measures, such as survival to adulthood and the age of first reproduction. We also reason that they may induce a long-term cumulative cost that may ultimately reduce longevity.

## Methods

### Study site, population under study, and sample collection

We used life-history and behavioural data, and 69 faecal samples collected non-invasively from 69 individually known female hyenas from 2010 to 2017. Individuals belonged to three clans respectively monitored on a nearly daily basis since 1987, 1989, and 1990, as part of an ongoing individual-based research project in the Serengeti National Park, Tanzania. Individuals were recognized by their distinctive patterns of spots, lesions, body scars, and other features such as ear-notches [57]. Maternal identity was determined by social behavioural interactions and was further confirmed by DNA microsatellite profiling [58]. Juvenile age was estimated by observing pelage development, body size, and movement coordination with an error of 7 days, or rarely, by direct observation of their date of birth [59]. Sex was determined at the age of approximately 3 months by the shape of external genitalia as described by [60]. Because females are philopatric and most adult males typically disperse from their natal clans when they reach adulthood [57,61], all the sampled individuals in this study were females.

The social rank(s) of the mothers of the sampled females was calculated using previously established methods [56]. Social ranks are determined from observation of submissive behaviours displayed in dyadic interactions recorded *ad libitum* during focal observations [62]. In each clan, submissive responses are used to construct a female linear dominance hierarchy following demographic and social changes (i.e. recruitment, deaths of adult females or coups) [56,54]. For comparison of rank positions across and within clans, we assigned standardised ranks, evenly distributing social ranks from the highest (standardised rank: +1) to the lowest rank (standardised rank: −1) within a clan (see [62]). Individuals younger than two years are assigned the same standardised ranks as the mothers raising [58]. In the case of adopted offspring we attributed the social rank of the surrogate mother and for jointly-raised offspring we used the rank of the mother with the highest value (n =4). The social rank(s) then were averaged from estimated date of birth to the first year of life of each individual.

As part of the long-term project, faeces are collected opportunistically, immediately after defecation. Following collection, they were stored in individual labelled bags and preserved in cool boxes with ice packs for no more than 3h after collection in the field until transport to the field station. In the field station, they were mechanically mixed and subsampled. Aliquots for faecal egg counts were preserved and stored in 4% formalin at room temperature, while the ones for measurements of faecal glucocorticoid metabolites (f-GCM), IgA (f-IgA) and mucin (f-mucin) were stored at -10°C, as well as during transport to Germany, where they were stored at -80°C [53]. Aliquots for DNA extraction were stored and transported in RNAlater at -10°C, transported to Germany on dry ice and finally stored at -80°C (Sigma-Aldrich, St. Louis, MO,USA).

We selected faecal samples collected within the first 6 months of life of the females, but when a female had no sample before this threshold, we used the sample collected at its earliest age (but before the age of 1 year, while they are still den-bound).

### Identification and quantification of gastrointestinal parasites (GIP)

We used a version of the McMaster egg flotation protocol for the detection (presence/absence) and counting of parasite eggs and oocysts [24]. We present the results here in faecal egg/oocyst counts per gram of faeces (FEC/FOC)*. Ancylostoma sp.* egg load was chosen as a predictor of performance measures in the models, based on the known negative effects of this parasite genus in this population [54,24,26]. To account for other parasites and the potential effect of polyparasitism, we used parasite richness (polyparasitism) as a measure of qualitative parasite diversity [24,26].

### Faecal immunological assays

Aliquots were lyophilized for 22 hours by a freeze-drier (Epsilon1-4 LSCplus, Martin Christ GmbH, Osterode, Germany), which was followed by a homogenization step with a mortar and pestle.

Our mucin assay was previously described in [26]. It can detect and quantify oligosaccharides released from mucin by differentiating O-linked and N-linked glycoproteins. Results are presented here as µmol oligosaccharide equivalents based on a standard curve created with N-acetylgalactosamine (Sigma-Aldrich(R)), Darmstadt, Germany). For positive controls we used Porcine stomach mucin (Sigma-Aldrich(R)), Darmstadt, Germany), and negative controls were blanks. All samples were run twice, and we only accepted results for which the coefficient of variation (CV) was below 5% and within the working range of the assay determined in [26].

For f-IgA detection and quantification we used a modified sandwich ELISA as described in [26]. We used saline extracts obtained from freeze-dried faecal samples with a protease-inhibitor MixM (Serva Electrophoresis GmbH, Heidelberg, Germany). The results are expressed as relative units (RU), as we used a pool of 72 wild hyena samples for the standard curve. We added negative controls, quality controls (in this case, hyena samples), and all samples were run twice to only accept results if the CV was below 5% and within the range established in [26].

### Faecal glucocorticoid metabolites (f-GCM)

Measurements of f-GCM were performed with an in-house competitive ELISA, validated for spotted hyenas [55]. The assay was performed with duplicates and controls, and we only accepted the results if the intra-assay CV was less than 10% and the inter-assay CV less than 15%, with the values fitting the curve of calibration [55]. The measurements are in ng/g. An outlier measurement of f-GCM was removed for all subsequent analyses (n=68).

### Statistical analysis

We conducted the analysis in R version 4.5.1 [63] and RStudio (version 2025.5.1.513 [64]). Tests were two-tailed with a level of significance set at 5%.

We investigated the effects of GIP, immune responses, allostatic load and maternal social rank (hereafter maternal rank) during early life on short- (age at first reproduction and survival to adulthood) and long-term (longevity) performance of 68 females. Following the definitions of Gicquel et al [48], survival to adulthood was used as a binary variable indicating whether the female survived to adulthood (at 730 days) or not. Age at first reproduction (AFR) was the age of a female when her first litter was born, regardless of offspring survival, in years. For the AFR model, as in Gicquel et al [48], we used a subset of the original dataset: all females that survived to at least adulthood were included, as well as those that had not yet reproduced but were still alive during our study period (n=48). Longevity was the female’s age when last observed alive, in years. For this last dataset, as in Ferreira et al [26], we included all the individuals from the original dataset.

To check if our results were affected by the use of an extended duration and broader definition of the early life period, we ran all models by including only samples collected before the 6 months of age (n = 51) as in Gicquel et al [48] and found similar results, (see Tables S4-S6).

We checked that the predictors were not strongly correlated with each other by calculating variance inflation factors with the R package Performance v. 0.14.0 [65]. Predictors were normalized for easier comparison of effects (rescaled to a mean of approximately 0 and a standard deviation of 1).

For survival to adulthood, we fitted a logistic regression model with survival (yes, no) as the response variable and maternal rank, *Ancylostoma sp.* egg load, polyparasitism, f-IgA, f-mucin and f-GCM, as continuous predictors.

To test the effects on AFR and longevity, we used Cox proportional hazards models to account for right-censored data using the packages Survival v. 3.8-3 [66] and Survminer v. 0.5.0 [67].

For AFR, as we were testing the effects on a subset of the original dataset (n=48), we chose the predictors maternal rank, *Ancylostoma sp.* faecal egg load and f-IgA levels given their significant influence in previous studies on parasite infections in juvenile Serengeti hyenas [24,26,48] with the inclusion of f-GCM - never explored before. For longevity we added again polyparasitism and f-mucin levels in the model.

Model assumptions for the logistic regression model were confirmed by using the package DHARMa v. 0.4.7 [68]. For the Cox-proportional-hazards models, we used the functions cox.zph from the package Survival v. 3.8-3 [66], ggcoxzph (i.e. Schoenfeld residuals) and ggcoxdiagnostics (i.e. outliers and Martingale test) both from the package Survminer v. 0.5.0 [67]. For longevity, because of severe violation of the proportional hazards assumption, one observation had to be removed from the dataset (n=67).

Log-likelihood ratio tests (LRT) were used to assess the significance of each predictor in the logistic regression model and for the Cox-proportional-hazards models by comparing them with a model with the predictor removed. Figures were produced with the packages ggeffects v. 2.3.0 [69] and ggplot2 v. 3.5.2 [70].

All the assays and protocols were run blind - no information regarding the identification of the individual, lab results or its life history and characteristics data was made available to the person running these assays or the statistical analysis.

## Results

We analysed the effects of GIP, local immune responses, allostatic load and maternal effects in the early-life stage on short and long-term performance of 68 female juveniles from 68 samples collected between 2010 and 2017 (Additional file 1: Fig. S1). The mean age at sample collection was 5 months (range: 1.40 - 10.20 months, 95% confident intervals (CI) 4.56 - 5.52 months), and 48 females survived to adulthood (71%). Among these 48 females, 35 gave birth to their first litter at an average age of 4 years (range 2.60 - 5.90 years, 95% CI 3.79 - 4.33 years). During the study period, 49 females died, and their mean longevity was 3.44 years (range 0.30 - 12 years, 95% CI 2.64 - 4.24 years).

We identified 8 different GIP taxa, three belonging to the phylum Nematoda, one to Apicomplexa and four to Platyhelminthes. *Ancylostoma sp.* was the most frequent GIP (91%, Table S1) and polyparasitism was observed in 91% of the individuals (Additional file 1: Fig. S2).

### Survival to adulthood is compromised by GIP infections and local immune responses

We found that survival to adulthood is positively influenced by maternal social rank and negatively by both *Ancylostoma sp.* egg loads and f-IgA levels (Figure 1, Table 1). Maternal rank had the highest contribution with an Odds Ratio (OR) of 5.82 (Table 1). In other words, when a mother’s rank goes up by just one unit, the offspring are nearly 6 times more likely to survive. Maternal rank is seen as having a protective effect and significantly improves an offspring’s probability of surviving to adulthood, although the precise magnitude of the effect is still not completely clear (95% CI [2.42-19.39]). The absolute gain in the survival probability is also not equal throughout the range of the maternal rank, being more pronounced within low to median rank levels (Figure 1 a). For each one-unit increase in *Ancylostoma sp.* egg load or f-IgA, the magnitude of the effect on the odds of surviving to adulthood is smaller and the direction is different (Table 1, Figure 1b and c). The presence of the *Ancylostoma sp.* eggs reduces the probability of surviving to adulthood - with an OR of 0.30, for each one-unit increase in *Ancylostoma sp.* eggs, an individual is 70% less likely to survive. The absolute gain in the survival probability is also not equal throughout the range for both *Ancylostoma sp.* egg load and f-IgA, being more pronounced within low to median levels (fig.1 b, c). *Ancylostoma sp.* egg load is seen making a higher contribution to this negative effect than f-IgA - where an unit increase on IgA levels leads to a decrease of the probability by 50% (Table 1).

**Figure 1.**
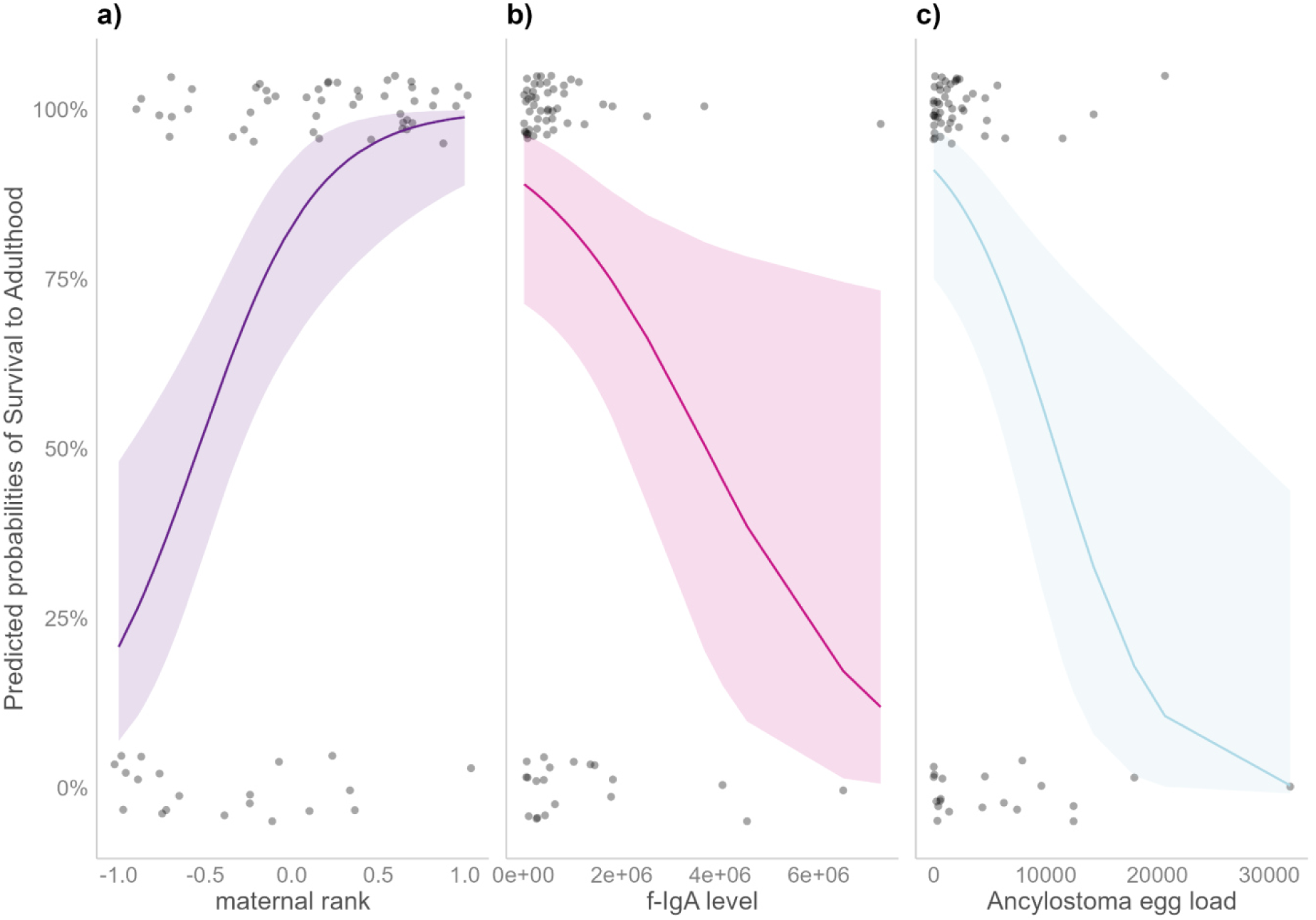
Survival to adulthood is influenced by maternal rank, f-IgA levels, and *Ancylostoma sp.* egg load. The bold lines indicate the predicted effect of maternal rank (a), f-IgA levels (b,) and *Ancylostoma sp.* egg load (c) on the probabilities of survival to adulthood in young female hyenas sampled before they were 12 months old. Shaded areas represent the 95% confidence intervals. Raw data of juveniles included in the model that survived (probability of survival to adulthood =1) and did not survive (probability of survival to adulthood =0) are shown as dots. The logistic regression model used for these predictions is presented in Table 1, n=68

**Table 1.** GIP, immunity, allostasis and maternal rank effects on the survival to adulthood of young female hyenas. We show the Odds Ratio (OR) from the full logistic regression model after variable normalization, with the corresponding 95% confidence interval (CI), including the results of the likelihood ratio test (LRT) and associated p-value, n=68. Estimates with * indicate predictors that have a significant effect in the survival of young hyenas sampled before they were 1 year old. - 95% CI does not include 1. CI: confident intervals. Polyparasitism is the number of other parasite taxa other than *Ancylostoma* sp. f-IgA, f-mucin and f-GCM are respectively markers of host local immunity and allostatic load.

|  | OR | 95% CI | LRT | p-value |
| --- | --- | --- | --- | --- |
| intercept | 4.96 | 2.18-15.52 | - |  |
| maternal rank* | 5.82 | 2.42-19.39 | 20.24 | p<0.001 |
| *Ancylostoma* sp. egg load* | 0.30 | 0.11-0.63 | 11.26 | p<0.001 |
| Polyparasitism | 1.24 | 0.65- 2.53 | 0.44 | 0.51 |
| f-IgA* | 0.46 | 0.21-0.91 | 4.84 | p<0.05 |
| f-mucin | 0.995 | 0.48-2.26 | 0.0001 | 0.99 |
| f-GCM | 0.63 | 0.24-1.19 | 1.98 | 0.16 |

Polyparasitism, f-mucin and fGCM did not influence female survival to adulthood (Table 1).

### Age at first reproduction (AFR) is not affected by the early life factors considered

For AFR, modelled here as chance of reproducing, we tested the effects of maternal rank, f-GCM levels, *Ancylostoma sp.* egg load, and the levels of f-IgA. We found that the probability to reproduce was not influenced by any of our predictors (Additional file 1:Table S2, Figure 2)..

**Figure 2.**
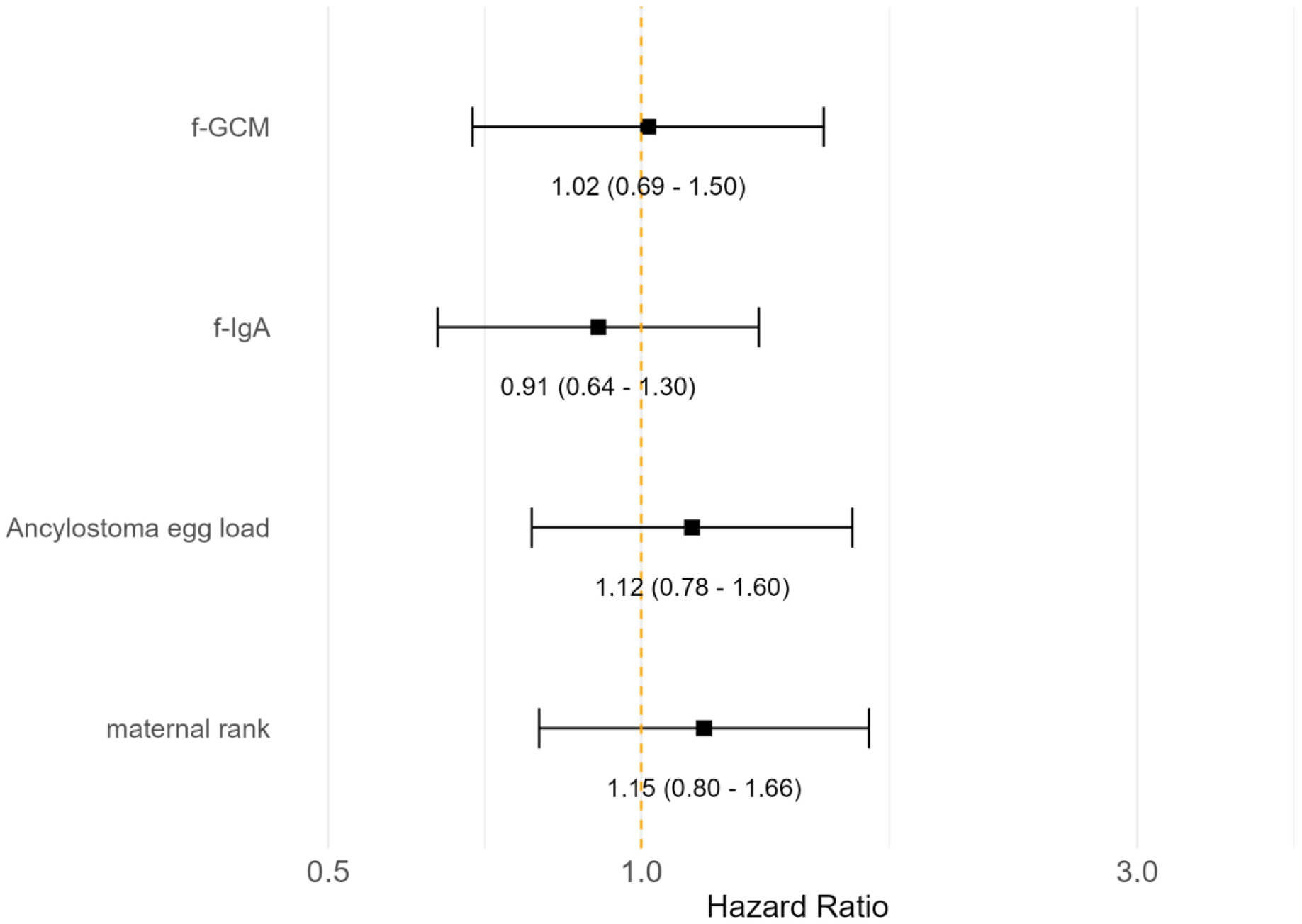
The effect of hazard ratios experienced by young female hyenas on AFR. Shown are the hazard ratios from the full Cox proportional hazard ratios model in Additional file 1:Table S2 in the age of first reproduction (AFR) of young hyenas sampled before they were 1 year old. Here we are assessing the chance of reproducing: a higher chance is indicative of an earlier age of first reproduction. Black squares with the corresponding 95% confidence intervals CI are represented as horizontal black lines, n=48. Predictors were rescaled for better visualization. Statistically significant predictors would be those whose CI do not cross the orange vertical interrupted line at OR =1 (p<0.05). f-IgA and f-GCM are respectively markers of host local immunity and allostatic load.

### Maternal social rank exerts a lasting effect on longevity

We tested the effects of maternal rank, f-GCM levels, *Ancylostoma sp.* egg load, polyparasitism, and the levels of f-IgA and f-mucin and found that the probability to die significantly decreased with increasing maternal rank (Hazard Ratio = 0.70, 95% CI= 0.52-0.96, LRT = 4.97, p<0.05) - a reduction of 30% in the hazard of dying is observed for each increase of one-unit (Additional file 1: table S3, figure 3). While holding constant all the other predictors to their median values, individuals with a low-ranking mother (−1 to 0) have a lower estimated survival probability and a higher risk of death at any given time compared with high-ranking mothers (0 to 1) (Fig.3 and 4).

**Figure 3.**
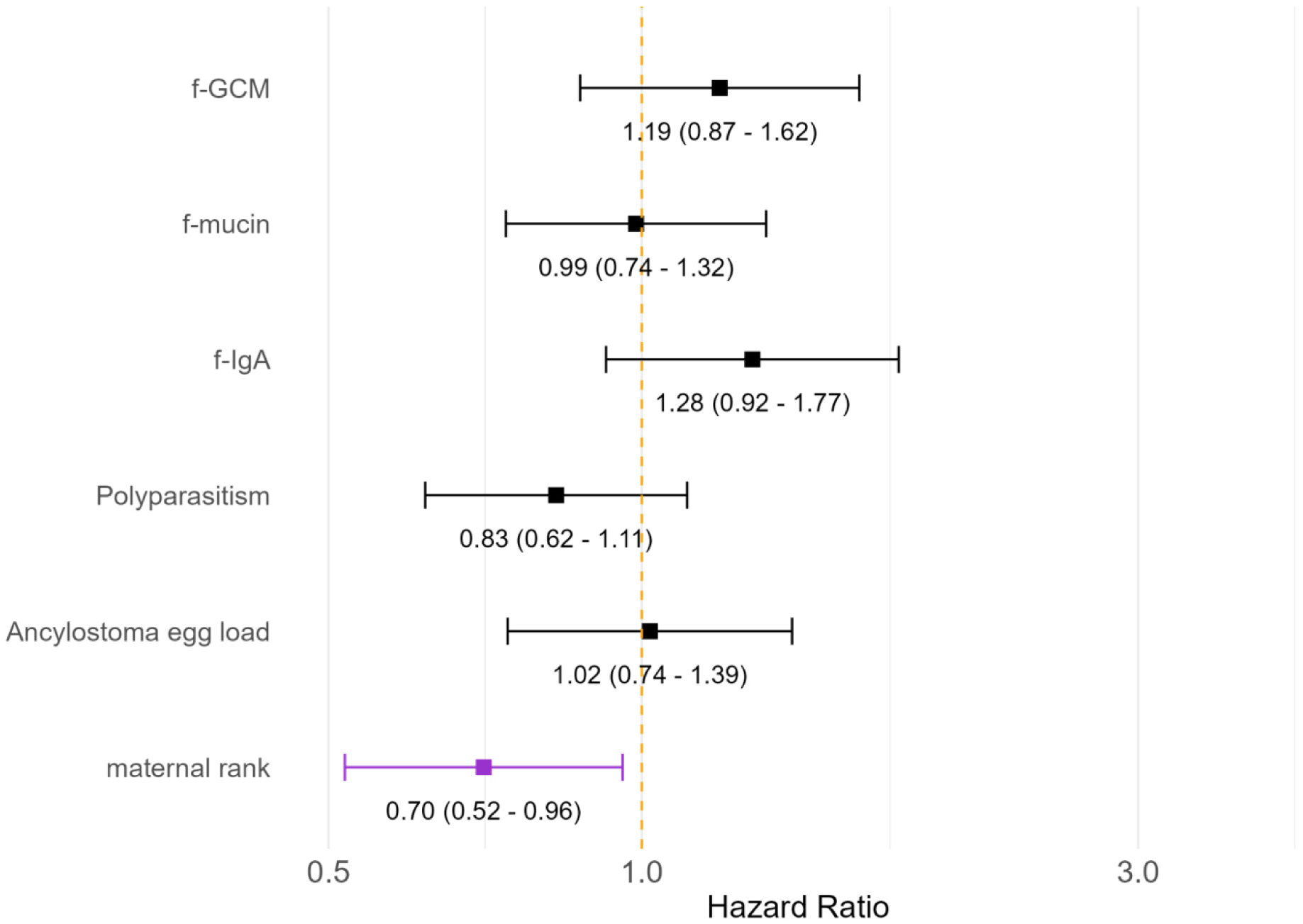
The effect of hazard ratios experienced by young female hyenas on longevity. Shown are the hazard ratios from the full Cox proportional hazard ratios model in Additional file 1:Table S3 in longevity of young hyenas sampled before they were 1 year old. Here we are assessing the chance of dying: a higher chance is indicative of an earlier death. Black squares with the corresponding 95% confidence intervals CI are represented as horizontal black lines, n=67. Predictors were rescaled for better visualization. Statistically significant predictors would be those whose CI do not cross the orange vertical interrupted line at OR =1 (p<0.05). Polyparasitism is the number of other parasite taxa other than *Ancylostoma* sp. f-IgA, f-mucin and f-GCM are respectively markers of host local immunity and allostatic load.

**Figure 4.**
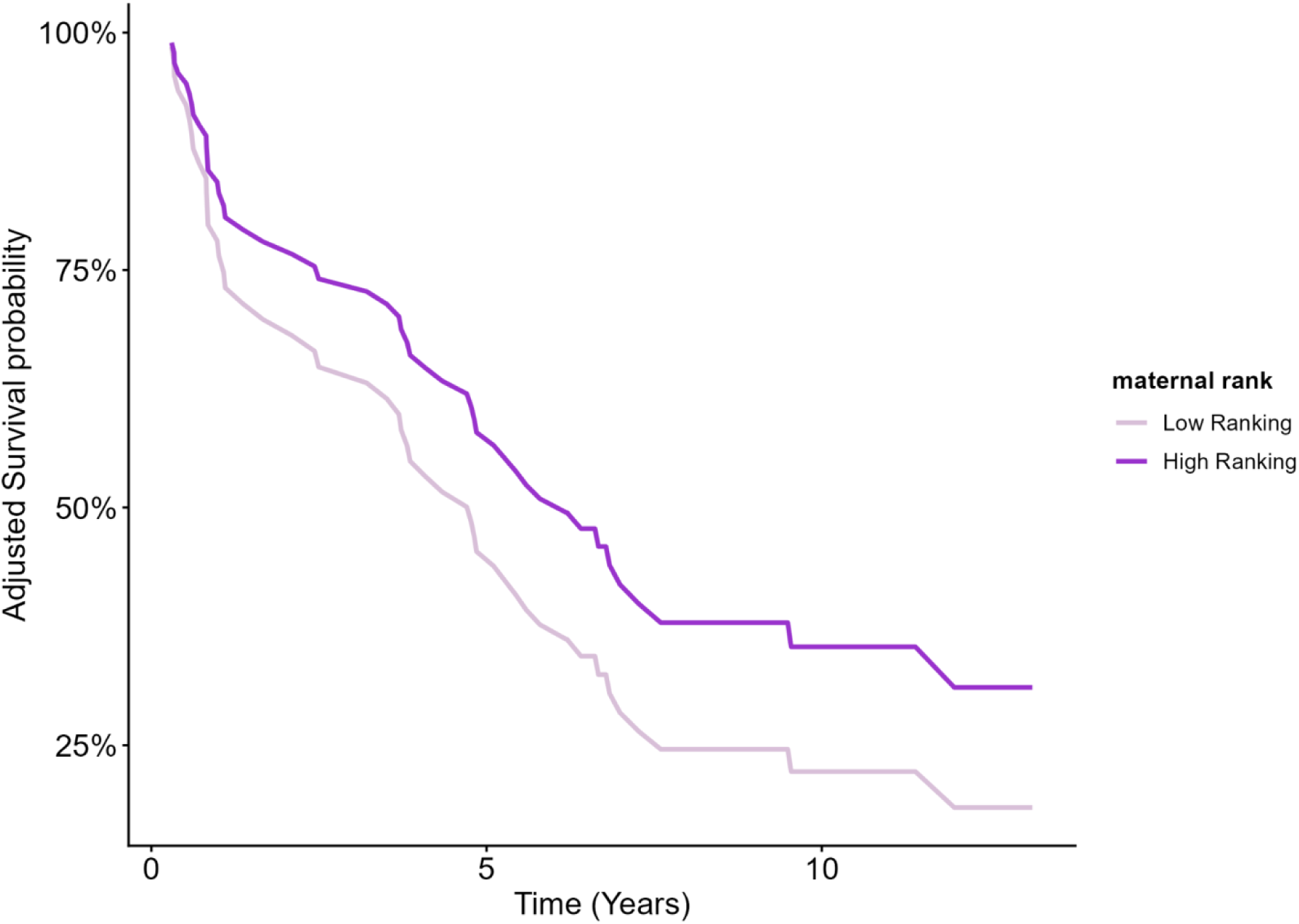
Longevity is influenced by maternal rank in young female hyenas. The bold lines show the adjusted survival curves indicating the estimated survival probability over time in years for two groups: those with a high-ranking mother 0 to 1 inclusive (dark purple) and those with a low-ranking mother, -1 to 0 (lighter purple) sampled when younger than 12 months of age.The full Cox proportional hazard model used for these predictions is presented in Additional file 1:table S3. All the remaining predictors are held constant at their median values, n=67.

All the other early life predictors had no effect (Additional file 1: Table S3, Figure 3).

## Discussion

We studied the effects of GIP, investment in local immunity and allostatic load during the early life of female spotted hyenas in the Serengeti on short and long-term measures of performance, while controlling for maternal social rank. For short-term effects, we found that the probability of surviving to adulthood was significantly reduced as *Ancylostoma sp.* egg loads and f-IgA concentration increased, suggesting that the potential damage and blood loss caused by the parasite and resource allocation trade-off between immunity and maintenance constrain survival. Conversely, maternal rank emerged as a strong positive predictor of both survival to adulthood and longevity of offspring. Age at first reproduction, however, appeared to be unaffected by any of the examined predictors. Our findings contribute novel insights to our understanding of how GIP, immunity, and allostatic load during early life shape the short and long-term performance of females of a social wild carnivore species.

In line with [24], our study reveals that high faecal egg loads of an energetically costly hookworm, *Ancylostoma sp.*, are associated with decrease in a short-term fitness metric: survival to adulthood. The established pathology of this hematophagous parasite in other mammalian species suggests a substantial potential for harm. *Ancylostoma sp*. attaches to the intestinal mucosa, inducing chronic blood loss through the release of anti-coagulating proteins, leading to anemia and nutrient deficiencies [71,72]. It also causes mucosal inflammation and tissue damage, resulting in retarded growth, weight loss, and severe gastrointestinal disturbances such as malabsorption, enteritis, and diarrhea, which can be fatal [71,72]. Secondary bacterial infections, arising from intestinal lining breaches, can further escalate to septicemia or peritonitis, increasing mortality risk [71,72]. Consistent with these broad pathological consequences, hookworm infections have been shown to negatively impact growth rates and contribute to mortality in wildlife populations, specifically in Australian sea lions (*Neophoca cinerea*), South American fur seals (*Arctocephalus australis*), and Northern fur seals (*Callorhinus ursinus*) [73–75]. Hookworms’ direct life cycle, environmental persistence (e.g. via paratenic hosts), and transmission routes - including potential transmammary and transplacental infection via larval reactivation during pregnancy and lactation [76,77,74] - may contribute to sustained transmission and reinfection thereby continuously impacting host performance and survival. In the Serengeti hyena population, lactating mothers exhibit higher infection rates and faecal *Ancylostoma sp*. egg loads than non-lactating females [54], potentially facilitating early-life infections and continuous maternal reinfection, thus perpetuating the parasite’s life cycle. Therefore, while we still cannot definitively establish a causal relationship, the consistently observed associations between *Ancylostoma sp.* egg load and survival coupled with the parasités known pathology and persistent life cycle strongly suggest a significant negative influence of *Ancylostoma sp.* infection during early life on short-term survival in hyenas. We did not observe a link between longevity and *Ancylostoma* sp. infection or immune responses during early life, contrary to previous studies which included both sexes [24,26]. This finding raises questions about potential sex and time-specific effects of early-life adversity that require further investigation to reconcile with existing emerging literature. For example, in wild ruminants, negative early life adversity effects were reported on male short-term fitness with factors like higher population sizes, harsh weather conditions and GIP having a greater influence [78]).

Parasite interactions can significantly influence fitness as well, whether through direct competition, environmental modification, or immune system modulation, often compounded by external environmental stressors [79,80]. While polyparasitism - the simultaneous presence of multiple parasite taxa - is known to detrimentally affect wildlife performance and fitness [15,43], our study’s use of simple parasite taxa presence as a metric likely limited our ability to detect these effects in our three models. This common and broad approach, encompassing diverse parasites with varied life cycles, niches and immunogenic potentials, does not capture the complexities of parasite co-occurrence, co-abundance, or their interactive effects (synergistic, additive, antagonistic, or independent). Furthermore, our approach overlooked factors influencing co-exposure, such as host age and spatiotemporal heterogeneity. Future research, employing refined polyparasitism metrics (e.g. accounting for individual parasite taxa effect), longitudinal sampling, and expanded datasets may reveal the potential effect of prior and subsequent infections, ultimately allowing the quantification of polyparasitism cumulative burdens [81].

We found that female survival to adulthood is also negatively associated with early-life f-IgA levels. Young mammals tend to be more susceptible to infections because their developing immune responses need to be trained, primed with exposure and require time to be effective [82,83]. This is particularly true for acquired immune responses, which differ from innate responses like mucin. IgAs are in between both branches [84,85], sharing features from innate and acquired immunity. Despite a potential for maternal lactogenic IgA transfer in mammals [86], we consistently observe a negative fitness effect associated with early life f-IgA and parasite burden in our study population (past studies [24,26]). This suggests that mounting an f-IgA-driven immune response is energetically costly, potentially diverting resources from growth and survival. It also raises the possibility that even if maternal IgA is transferred to hyena juveniles through lactation, the amount received might not fully offset the energetic demands of their own antibody-mediated immune responses. Furthermore, considering our previous findings on the diverse associations between immune measures and gut microbiome components [87], our current results could suggest a possible life-compromising dysregulation of the gut microbiome. Future research focusing on pathogen-specific Ig levels on this life stage will be crucial to clarify the reason behind this survival compromise.

Contrary to our expectation, we did not find an effect of f-GCM on the measures of performance considered. Previous research in free-ranging spotted hyenas has already identified relationships between life history stages (e.g. lactation), behaviour (e.g.sibling rivalry), and allostatic load [62,88]. Our attempt to fill the knowledge gap on the effect of f-GCM on performance in this species shows that this discrepancy may reveal the complex, context-dependent nature of allostatic load effects [89–91], as evidenced by the variable associations between allostatic load and GIP mainly found in wildlife studies, involving experimental parasite reduction [92]. These associations are highly dependent on the specific model under investigation, encompassing parasite [93], host (e.g. social avoidance behaviour [94,95]), immune response (e.g. f-IgA [96]) and environmental factors [97]. These are embedded within complex interactions between hosts (i.e. tolerance), resource availability, parasitic infection and reproduction [98,97,99]. In our study, the same may have attenuated the expected f-GCM’s impact on performance. Alternatively, f-GCM, while providing an integrated measure of adrenal activity over an extended period of time [100], may not fully capture the relevant allostatic pathways and proxies in action. A more comprehensive, non-invasive, species-specific allostatic load index using multiple biomarkers (neuroendocrine, cardiovascular, immune, etc.) and temporal dynamics might reveal obscured relationships by our single measurement [101]. Furthermore, the inherent individual stability of fitness parameters compared to the dynamic and large fluctuations of f-GCM [89] are likely to have limited our predictive power when we resort to a single time-point sampling, potentially underrepresenting its effect or masking both acute and chronic stress effects. For instance, acute increases in glucocorticoids can temporarily enhance immune response, whereas chronic elevations may lead to immunomodulation or even suppression [37,102].

Consistent with previous research on free-ranging hyenas [56,24,26,51,48], we found that maternal rank positively influenced offspring survival to adulthood and longevity, highlighting the crucial role of maternal effect in early development, a pattern observed across mammals [103]. In Serengeti hyenas, maternal rank affects a mother’s access to resources, foraging effort in terms of commuting behaviour, milk transfer and within-litter social dynamics, directly influencing cub growth rate, survival probability and age at first reproduction [59,56,46]. Maternal rank also potentially influences parasite-host interactions. Low-ranking lactating females exhibit higher *Ancylostoma sp*. egg loads and allostatic load than non-lactating females [62,54], potentially contributing to increased cub infection rates [24]. Furthermore, suboptimal milk supply and possibly reduced passive immunity transfer during the critical first six months of life, coupled with increased parasite transmission, can compromise cub development and contribute to high parasite loads [24,26]. Such factors probably hinder a hyena’s ability to allocate resources effectively, impacting their capacity to manage challenges such as GIP during growth and beyond. Therefore, maternal rank likely serves as a complex composite index, reflecting maternal investment and experiences, cub nutrition, parasitism, and immunity, which profoundly shape short and long-term life outcomes.

Our study found no effect of any of our predictors on the age at first reproduction (AFR), our measure of reproductive performance at the start of female reproductive career. In other species, AFR is influenced by various factors, including parasitism, body condition, and social status [104–106]. In a previous study that considered the impact of several predictors aimed at capturing variation in ecological, social, and demographic conditions in more than 600 juvenile female spotted hyenas, only maternal rank influenced the AFR [48]. The absence of an effect of GIP on AFR in our study may be due to our comparatively smaller sample size and several explanations. Firstly, parasites, though known to affect reproduction, may not have reached a threshold high enough to impact AFR in our population, or tolerance mechanisms to reduce their negative effect might be at play [107]. Secondly, body condition, a strong AFR predictor in many species [108,109], was not directly measured in our study, and our proxies might not adequately represent it (see [110]). Thirdly, social factors, like maternal rank, previously linked to AFR in the Serengeti hyena population [56,48], might have been obscured by our smaller sample size, potentially biased subset towards individuals that were alive and for whom samples could be collected for analysis. Finally, the lack of f-GCM influence on AFR suggests early-life stress mediation by this pathway might not be a significant AFR determinant in female spotted hyenas. It is also possible that the sampled individuals had time to recover from the potential energetic deficits and damage by the time they can reproduce, and these infections are not entirely detrimental, playing a beneficial selection role - a hypothesis requiring further investigation. Ultimately, AFR is likely a product of complex interactions across life stages, and our single-time point measurements during the early life period may fail to capture these dynamics [111].

## Conclusion

Our study on an unmanaged wild Serengeti hyena population integrates and determines how early life GIP, immunity, and maternal effects uniquely shape both short- and long-term fitness measures of females. Our non-invasive approach, which also avoided antiparasitic drugs, eliminated concerns about uncontrolled disturbance effects, drug efficacy [112], or resistance, contributing to the slowly growing number of these wildlife studies that aim to understand the full scope of parasite-host dynamics in wild, unmanipulated populations. Despite a modest sample size, our results show a significant negative impact of GIP and immunity on juvenile survival to adulthood, emphasizing the critical cost of early-life parasitism. However, the costs of early infection do not necessarily carry over into early adulthood or influence investment in reproduction, as neither measure affects longevity or the AFR. Further studies with larger sample sizes and repeated measures are needed to fully evaluate these effects, potential confounding variables and alternative mechanistic pathways. Our research thus improves our knowledge of the intricate and complex dynamic associations between maternal influence, GIP, immunity, and allostatic load during early life. Collectively, our results confirm the enduring significance of maternal investment and corroborates the established paradigm across various species that early-life experiences can exert critical and lasting effects on fitness trajectories.

## Acknowledgments

We thank the Tanzania Commission for Science and Technology (COSTECH), the Tanzanian National Parks Authority (TANAPA) and Tanzanian Wildlife Research Institute (TAWIRI) for supporting the fieldwork: grating the research permits and necessary permissions.

We thank D. Thierer, K. Pohle, F. Webster, C. Bost, J. Dalijono, J. Hens, E. Böhm, M. Albrecht and K. Grassal for lab assistance. We thank A. Francis, T. Shabani, M. Andris, N. Boyer, T. Golla, K. Goller, N. Gusset-Burgener, B. Kostka, M. Lindson, D. Thierer, A. Türk and K. Wilhelm for field and technical assistance. We also thank Prof. Dr. Friedrieke Ebner for her valuable inputs during the past thesis advisory committee meetings.

## Availability of data and materials

The dataset and code supporting the conclusions of this article are available in Zenodo repository, (https://doi.org/10.5281/zenodo.17957004).

## Author’s contributions

SPVS, SCMF, MG and SB conceptualised the study. MLE, GC and HH acquired funding. SCMF, SM, MLE, HH and SB conducted fieldwork. SPSV, MMV and SCMF performed the laboratory work for faecal egg counts and immune measures. SM, SK and JW performed laboratory work for faecal glucocorticoid metabolite measures. SPVS analysed the data. SPVS wrote the manuscript. SPVS created the graphical abstract. All authors contributed to the review and editing of the manuscript.

## Ethics approval

The fieldwork was authorized, supported and research permits were issued by the Tanzania Commission for Science and Technology (COSTECH), the Tanzanian National Parks Authority (TANAPA), the Tanzanian Wildlife Research Institute (TAWIRI). Lab procedures and protocols were developed and implemented under the evaluation of the Leibniz Institute for Zoo and Wildlife Research Ethics Committee on Animal Welfare (permit number: 2017-11-02). We complied with all ethical regulations and our samples were exported with all the necessary permits.

## Consent for publication

All the authors consent for publication.

## Supplementary information

### Additional file 1

#### Sampled individuals

**Fig. S1.**
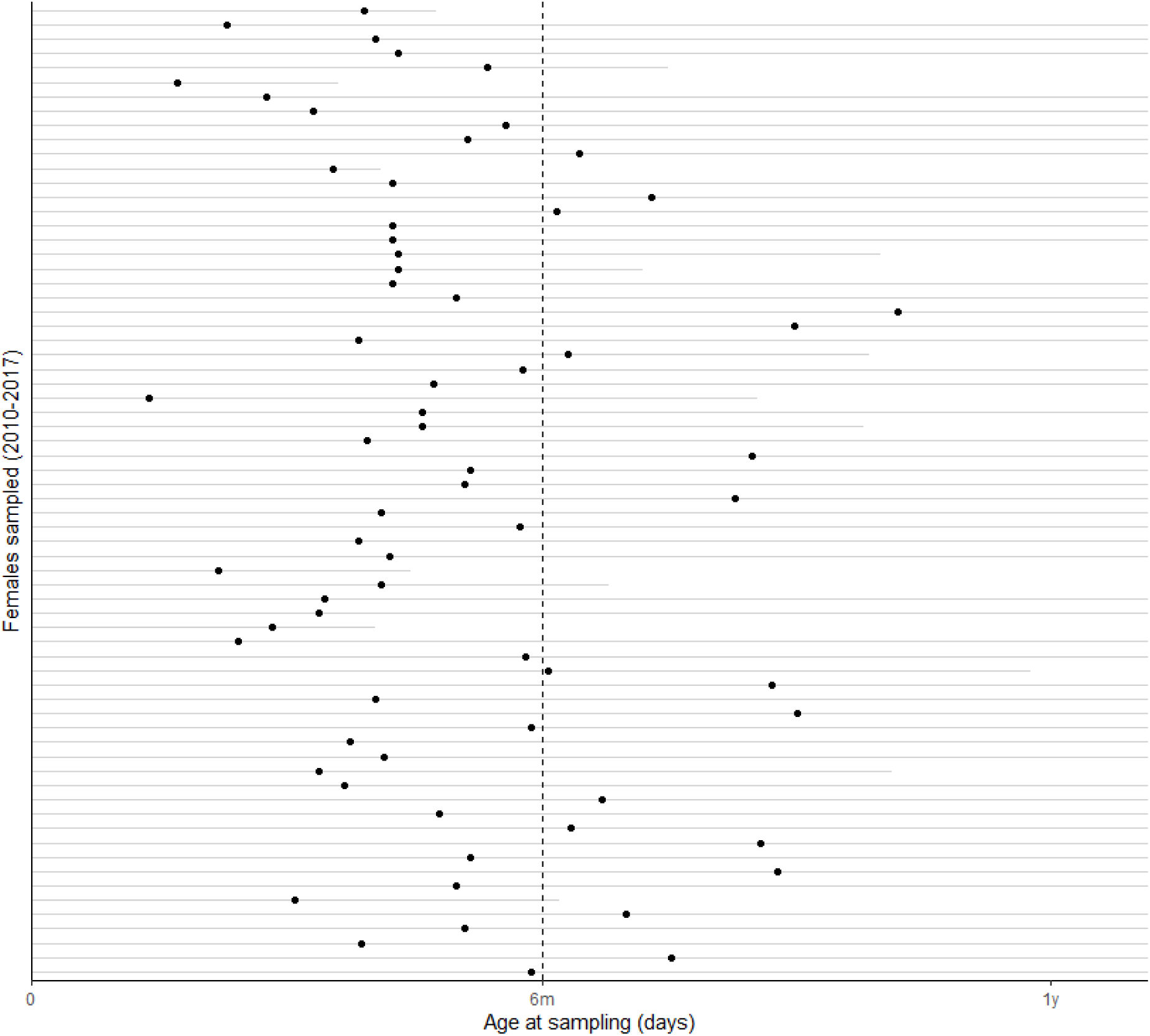
Female lifelines and moments when they were sampled for all models (2010-2017) n=68. Each horizontal line represents one female. Showcased are the moments when they were sampled (full black dots) and when they died (line ending). Represented by a dashed vertical line is the moment when they were 6 months old (confirmation threshold).

#### Parasitism quantification measures

**Table S1.** Parasites detected and quantified in 68 samples: prevalence (%), mean intensity-arithmetic mean of the number of individual parasites of a particular species per infected host in a sample, median intensity - indice of parasite distribution in infected hosts, and mean abundance -arithmetic mean of the number of individual parasites of a particular species per examined host including both infected or not, in accordance with [113], n=68. ** polyparasitism does not include *Ancylostoma* sp.

| Parasites | Phila | Prevalence | Mean Intensity | Median Intensity | Mean abundance |
| --- | --- | --- | --- | --- | --- |
| *Ancylostoma* | Nematoda | 91% | 3764.28 | 1487.5 | 3432.14 |
| *Cystoisospora* | Apicomplexa | 59% | 3831.56 | 200 | 2253.86 |
| *Diphyllobothrium* | Platyhelminthes | 44% | 21202.08 | 6100 | 9353.86 |
| *Dipylidium* | Platyhelminthes | 38% | - | - | - |
| *Trichuris* | Nematoda | 6% | 31.25 | 25 | 25 |
| Taeniidae | Platyhelminthes | 12% | 28.13 | 25 | 3.31 |
| Spirurida | Nematoda | 9% | 308.33 | 87.5 | 27.21 |
| *Mesocistoides* | Platyhelminthes | 34% | 79.89 | 50 | 27.02 |
| *Polyparasitism* ** | - | 91% | - | - | - |

#### Study sampling design

**Fig. S2.**
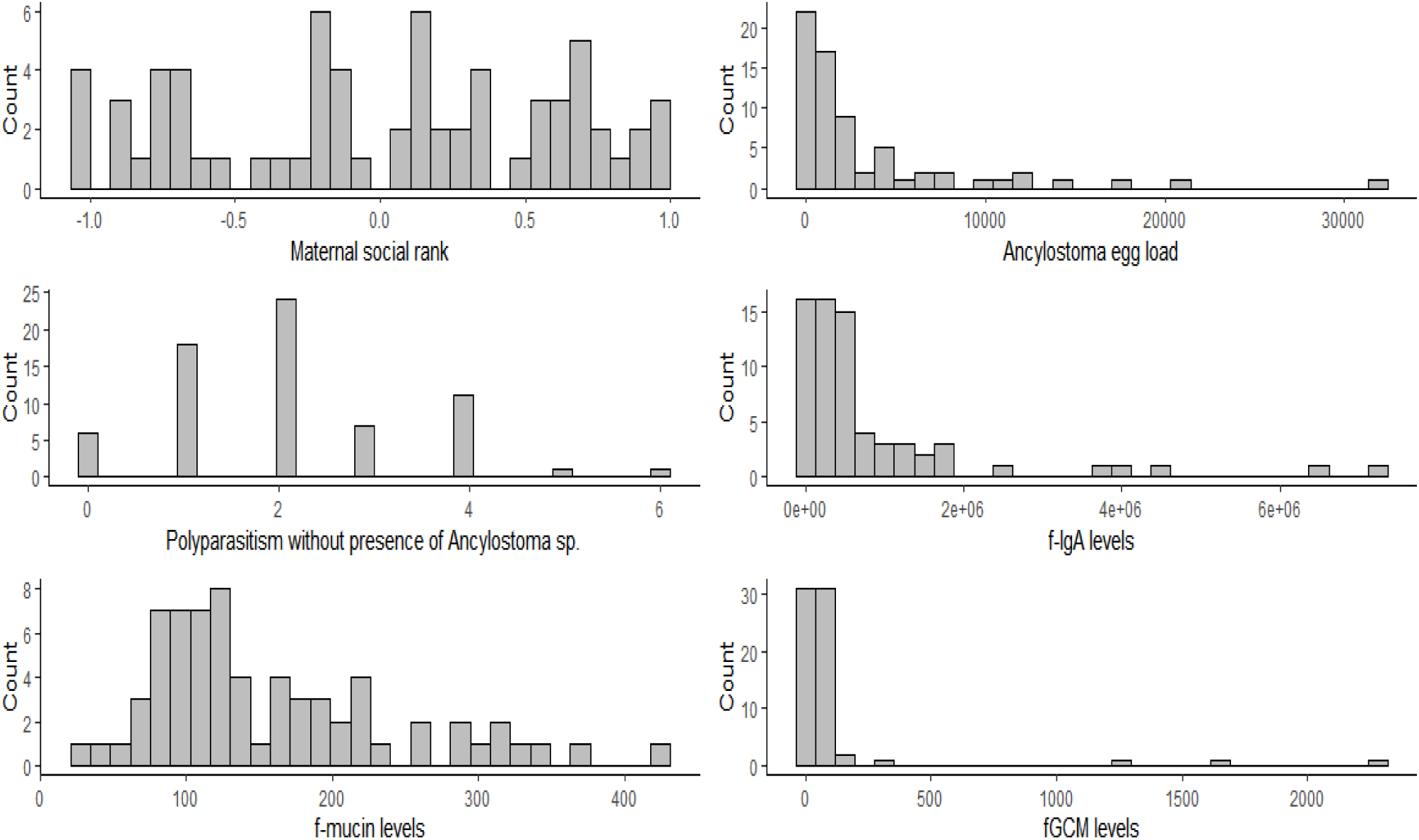
Sampling design of the study. A) Histogram of averaged individual maternal standardised social rank across early life of sampled juveniles. B) Histogram of individual counted feacal egg of *Ancylostoma* sp. across samples. C) Frequency count of sampled individuals with more than one GIP, excluding presence of *Ancylostoma* sp. D) Histogram of the individual f-IgA levels measured across samples. E) Histogram of the individual f-mucin levels measured across samples. F) Histogram of the individual f-GCM levels measured across samples. Individual count includes 68 individuals and sample count includes 68 samples.

### Early life (less than 1 year old of age)

#### Age at first reproduction (AFR) is not affected by the early life factors considered

**Table S2.** GIP, immunity, allostasis and maternal rank effects on AFR of young female hyenas. We show the Hazard Ratios (HR) from the full Cox-proportional-hazards model, with the corresponding 95% Confidence Intervals (CI), including the results of the LRT test and associated p-value, n=48. Estimates with * indicate significant predictors in AFR (chance of reproducing) of young hyenas sampled before they were 1 year old - 95% CI does not include 1. f-IgA and f-GCM are respectively markers of host local immunity and allostatic load.

|  | HR | 95%CI | LRT | p-value |
| --- | --- | --- | --- | --- |
| maternal rank | 1.15 | 0.80 -1.66 | 0.56 | 0.45 |
| *Ancylostoma* sp. egg load | 1.12 | 0.78 -1.60 | 0.34 | 0.56 |
| f-IgA | 0.91 | 0.64 - 1.30 | 0.30 | 0.59 |
| f-GCM | 1.02 | 0.69 -1.50 | 0.01 | 0.94 |

#### Maternal social rank exerts a lasting effect on longevity

**Table S3.** GIP, immunity, allostasis and maternal rank effects on longevity of young female hyenas. We show the Hazard Ratios (HR) from the full Cox proportional hazards model, with the corresponding 95% Confidence Intervals (CI), including the results of the LRT test and associated p-value, n=67. Estimates with * indicate significant predictors in longevity (chance of dying) of young hyenas sampled before they were 1 year old - 95% CI does not include 1. Polyparasitism is the number of other parasite taxa other than *Ancylostoma* sp. f-IgA, f-mucin and f-GCM are respectively markers of host local immunity and allostatic load.

| Predictors | HR | 95%CI | LRT | p-value |
| --- | --- | --- | --- | --- |
| maternal rank* | 0.70 | 0.52 - 0.96 | 4.97 | <0.05 |
| *Ancylostoma* sp. egg load | 1.02 | 0.74 - 1.39 | 0.01 | 0.91 |
| Polyparasitism | 0.83 | 0.62 - 1.11 | 1.73 | 0.19 |
| f-IgA | 1.28 | 0.92 - 1.77 | 1.95 | 0.16 |
| f-mucin | 0.99 | 0.74 - 1.32 | 0.01 | 0.93 |
| f-GCM | 1.19 | 0.87 - 1.62 | 1.02 | 0.31 |

### Early life (confirmation for before 6 months of age)

#### Survival to adulthood is compromised by GIP infections and local immune responses

**Table S4.** GIP, immunity, allostasis and maternal rank effects on survival to adulthood of young female hyenas. We show the Odds Ratios (OR) from the full logistic regression model after normalisation, with the corresponding 95% Confidence Intervals (CI), including the results of the likelihood ratio test (LRT) and associated p-value, n=51. Estimates with * indicate predictors that have a significant effect in the survival of young hyenas sampled before they were 6 months old - 95% CI does not include 1. CI: confidence intervals. Polyparasistism is the number of the other parasite taxa other than *Ancylosotoma* sp. f-IgA, f-mucin and f-GCM are respectively markers of host local immunity and allostatic load.

| Predictors | OR | 95%CI | LRT | p-value |
| --- | --- | --- | --- | --- |
| intercept | 3.50 | 1.52-10.56 |  |  |
| maternal rank* | 5.68 | 2.21-20.62 | 16.19 | p<0.001 |
| *Ancylostoma* sp. egg load* | 0.31 | 0.09-0.72 | 7.95 | p<0.01 |
| Polyparasitism | 1.20 | 0.51- 3.08 | 0.17 | 0.68 |
| f-IgA* | 0.37 | 0.13 -0.88 | 5.07 | p<0.05 |
| f-mucin | 1.01 | 0.43- 2.87 | 0.0002 | 0.99 |
| f-GCM | 1.01 | 0.37- 2.30 | 0.0005 | 0.98 |

#### Age at first reproduction (AFR) is not affected by the early life factors considered

**Table S5.** GIP, immunity, allostasis and maternal rank effects on AFR of young female hyenas. We show the Hazard Ratios (HR) from the full Cox proportional hazards model, with the corresponding 95% Confidence Intervals (CI), including the results of the LRT test and associated p-value, n=34. Estimates with * indicate significant predictors - 95% CI does not include 1- in the AFR (chance of reproducing) of young female hyenas sampled before they were 6 months old. f-IgA, f-mucin and f-GCM are respectively markers of host local immunity and allostatic load.

| Predictors | HR | 95%CI | LRT | p-value |
| --- | --- | --- | --- | --- |
| maternal rank | 1.10 | 0.68 - 1.76 | 0.15 | 0.70 |
| *Ancylostoma* sp. egg load | 1.19 | 0.78 - 1.81 | 0.55 | 0.46 |
| f-IgA | 0.94 | 0.61 - 1.46 | 0.07 | 0.78 |
| f-GCM | 1.01 | 0.65 - 1.59 | 0.003 | 0.96 |

#### Maternal social rank exerts a lasting effect on longevity

**Table S6.** GIP, immunity, allostasis and maternal rank effects on longevity of young female hyenas. We show the Hazard Ratios (HR) from the full Cox proportional hazards model, with the corresponding 95% Confidence Intervals (CI), including the results of the LRT test and associated p-value, n=51. Estimates with * indicate significant predictors - 95% CI does not include 1- in longevity (chance of dying) of young hyenas sampled before they were 6 months old. Polyparasitism is the number of other parasite taxa other than *Ancylostoma* sp, f-IgA, f-mucin and f-GCM are respectively markers of host local immunity and allostatic load.

| Predictors | HR | 95%CI | LRT | p-value |
| --- | --- | --- | --- | --- |
| maternal rank* | 0.69 | 0.48-1.01 | 3.83 | 0.05 |
| *Ancylostoma* sp. egg load | 1.27 | 0.84-1.93 | 1.21 | 0.27 |
| polyparasitism | 0.94 | 0.67-1.32 | 0.11 | 0.74 |
| f-IgA | 1.48 | 0.97-2.26 | 2.82 | 0.09 |
| f-mucin | 0.88 | 0.59-1.30 | 0.47 | 0.49 |
| f-GCM | 1.02 | 0.72-1.45 | 0.01 | 0.92 |

## References

1. Lindström J. Early development and fitness in birds and mammals. Trends Ecol Evol. 1999 Sept;14(9):343–8.

2. Dettmer AM, Chusyd DE. Early life adversities and lifelong health outcomes: A review of the literature on large, social, long-lived nonhuman mammals. Neurosci Biobehav Rev. 2023 Sept;152:105297.

3. Descamps S, Boutin S, Berteaux D, McAdam AG, Gaillard J. Cohort effects in red squirrels: the influence of density, food abundance and temperature on future survival and reproductive success. J Anim Ecol. 2008 Mar;77(2):305–14.

4. Marshall HH, Vitikainen EIK, Mwanguhya F, Businge R, Kyabulima S, Hares MC, et al. Lifetime fitness consequences of early-life ecological hardship in a wild mammal population. Ecol Evol. 2017 Mar;7(6):1712–24.

5. Charpentier MJE, Tung J, Altmann J, Alberts SC. Age at maturity in wild baboons: genetic, environmental and demographic influences. Mol Ecol. 2008 Apr;17(8):2026–40.

6. Berger V, Lemaître JF, Allainé D, Gaillard JM, Cohas A. Early and adult social environments have independent effects on individual fitness in a social vertebrate. Proc R Soc B Biol Sci. 2015 Aug 22;282(1813):20151167.

7. Monaghan P. Early growth conditions, phenotypic development and environmental change. Philos Trans R Soc Lond B Biol Sci. 2008 May 12;363(1497):1635–45.

8. Duffy KA, McLaughlin KA, Green PA. Early life adversity and health-risk behaviors: proposed psychological and neural mechanisms. Ann N Y Acad Sci. 2018 Sept;1428(1):151–69.

9. Langenhof MR, Komdeur J. Why and how the early-life environment affects development of coping behaviours. Behav Ecol Sociobiol. 2018 Mar;72(3):34.

10. Hotez PJ. Empowering Women and Improving Female Reproductive Health through Control of Neglected Tropical Diseases. PLoS Negl Trop Dis. 2009 Nov 24;3(11):e559.

11. Yeshanew S, Bekana T, Truneh Z, Tadege M, Abich E, Dessie H. Soil-transmitted helminthiasis and undernutrition among schoolchildren in Mettu town, Southwest Ethiopia. Sci Rep. 2022 Mar 7;12(1):3614.

12. Perri AF, Mejía ME, Licoff N, Lazaro L, Miglierina M, Ornstein A, et al. Gastrointestinal parasites presence during the peripartum decreases total milk production in grazing dairy Holstein cows. Vet Parasitol. 2011 June;178(3–4):311–8.

13. Borkowski EA, Avula J, Redman EM, Sears W, Lillie BN, Karrow NA, et al. Correlation of subclinical gastrointestinal nematode parasitism with growth and reproductive performance in ewe lambs in Ontario. Prev Vet Med. 2020 Dec;185:105175.

14. Acerini CI, Morris S, Morris A, Kenyon F, McBean D, Pemberton JM, et al. Helminth parasites are associated with reduced survival probability in young red deer. Parasitology. 2022 Nov;149(13):1702–8.

15. Alzaga V, Vicente J, Villanua D, Acevedo P, Casas F, Gortazar C. Body condition and parasite intensity correlates with escape capacity in Iberian hares (Lepus granatensis). Behav Ecol Sociobiol. 2008 Mar;62(5):769–75.

16. Albon SD, Stien A, Irvine RJ, Langvatn R, Ropstad E, Halvorsen O. The role of parasites in the dynamics of a reindeer population. Proc R Soc Lond B Biol Sci. 2002 Aug 7;269(1500):1625–32.

17. Hillegass MA, Waterman JM, Roth JD. Parasite removal increases reproductive success in a social African ground squirrel. Behav Ecol. 2010 July 1;21(4):696–700.

18. Stien A, Irvine RJ, Ropstad E, Halvorsen O, Langvatn R, Albon SD. The impact of gastrointestinal nematodes on wild reindeer: experimental and cross-sectional studies. J Anim Ecol. 2002 Nov;71(6):937–45.

19. Newey S, Thirgood S. Parasite–mediated reduction in fecundity of mountain hares. Proc R Soc Lond B Biol Sci [Internet]. 2004 Dec 7 [cited 2025 Jan 9];271(suppl_6). Available from: https://royalsocietypublishing.org/doi/10.1098/rsbl.2004.0202

20. Shanebeck KM, Besson AA, Lagrue C, Green SJ. The energetic costs of sub-lethal helminth parasites in mammals: a meta-analysis. Biol Rev. 2022 Oct;97(5):1886–907.

21. Ardia DR, Parmentier HK, Vogel LA. The role of constraints and limitation in driving individual variation in immune response. Funct Ecol. 2011 Feb;25(1):61–73.

22. Watson RL, McNeilly TN, Watt KA, Pemberton JM, Pilkington JG, Waterfall M, et al. Cellular and humoral immunity in a wild mammal: Variation with age & sex and association with overwinter survival. Ecol Evol. 2016 Dec;6(24):8695–705.

23. Benavides JA, Huchard E, Pettorelli N, King AJ, Brown ME, Archer CE, et al. From parasite encounter to infection: Multiple-scale drivers of parasite richness in a wild social primate population. Am J Phys Anthropol. 2012 Jan;147(1):52–63.

24. Ferreira SCM, Hofer H, Madeira De Carvalho L, East ML. Parasite infections in a social carnivore: Evidence of their fitness consequences and factors modulating infection load. Ecol Evol. 2019 Aug;9(15):8783–99.

25. Gesquiere LR, Habig B, Hansen C, Li A, Freid K, Learn NH, et al. Noninvasive measurement of mucosal immunity in a free-ranging baboon population. Am J Primatol. 2020 Feb;82(2):e23093.

26. Ferreira SCM, Veiga MM, Hofer H, East ML, Czirják GÁ. Noninvasively measured immune responses reflect current parasite infections in a wild carnivore and are linked to longevity. Ecol Evol. 2021 June;11(12):7685–99.

27. Pelaseyed T, Bergström JH, Gustafsson JK, Ermund A, Birchenough GMH, Schütte A, et al. The mucus and mucins of the goblet cells and enterocytes provide the first defense line of the gastrointestinal tract and interact with the immune system. Immunol Rev. 2014 July;260(1):8–20.

28. Hansson GC. Mucins and the Microbiome. Annu Rev Biochem. 2020 June 20;89(1):769–93.

29. Mantis NJ, Rol N, Corthésy B. Secretory IgA’s complex roles in immunity and mucosal homeostasis in the gut. Mucosal Immunol. 2011 Nov;4(6):603–11.

30. Pabst O, Izcue A. Secretory IgA: controlling the gut microbiota. Nat Rev Gastroenterol Hepatol. 2022 Mar;19(3):149–50.

31. Sheldon BC, Verhulst S. Ecological immunology: costly parasite defences and trade-offs in evolutionary ecology. Trends Ecol Evol. 1996 Aug;11(8):317–21.

32. Lochmiller RL, Deerenberg C. Trade-offs in evolutionary immunology: just what is the cost of immunity? Oikos. 2000 Jan;88(1):87–98.

33. Romero LM, Dickens MJ, Cyr NE. The reactive scope model — A new model integrating homeostasis, allostasis, and stress. Horm Behav. 2009 Mar;55(3):375–89.

34. McEwen BS, Wingfield JC. What is in a name? Integrating homeostasis, allostasis and stress. Horm Behav. 2010 Feb;57(2):105–11.

35. Landys MM, Ramenofsky M, Wingfield JC. Actions of glucocorticoids at a seasonal baseline as compared to stress-related levels in the regulation of periodic life processes. Gen Comp Endocrinol. 2006 Sept;148(2):132–49.

36. MacDougall-Shackleton SA, Bonier F, Romero LM, Moore IT. Glucocorticoids and “Stress” Are Not Synonymous. Integr Org Biol. 2019 Jan 1;1(1):obz017.

37. Sapolsky RM, Romero LM, Munck AU. How Do Glucocorticoids Influence Stress Responses? Integrating Permissive, Suppressive, Stimulatory, and Preparative Actions*. Endocr Rev. 2000 Feb 1;21(1):55–89.

38. Cabezas S, Blas J, Marchant TA, Moreno S. Physiological stress levels predict survival probabilities in wild rabbits. Horm Behav. 2007 Mar;51(3):313–20.

39. Sheriff MJ, Krebs CJ, Boonstra R. The sensitive hare: sublethal effects of predator stress on reproduction in snowshoe hares. J Anim Ecol. 2009 Nov;78(6):1249–58.

40. Benedek I, Altbӓcker V, Molnár T. Stress reactivity near birth affects nest building timing and offspring number and survival in the European rabbit (Oryctolagus cuniculus). Cameron EZ, editor. PLOS ONE. 2021 Jan 29;16(1):e0246258.

41. Murray CM, Stanton MA, Wellens KR, Santymire RM, Heintz MR, Lonsdorf EV. Maternal effects on offspring stress physiology in wild chimpanzees. Am J Primatol. 2018 Jan;80(1):e22525.

42. Anzà S, Heistermann M, Ostner J, Schülke O. Early prenatal but not postnatal glucocorticoid exposure is associated with enhanced HPA axis activity into adulthood in a wild primate. Proc R Soc B Biol Sci. 2025 Jan;292(2039):20242418.

43. Akinyi MY, Jansen D, Habig B, Gesquiere LR, Alberts SC, Archie EA. Costs and drivers of helminth parasite infection in wild female baboons. Cattadori I, editor. J Anim Ecol. 2019 July;88(7):1029–43.

44. Matthews, L. H. Reproduction in the spotted hyaena, *Crocuta crocuta* (erxleben). Philos Trans R Soc Lond B Biol Sci. 1939 July 5;230(565):1–78.

45. Holekamp KE, Cooper SM, Katona CI, Berry NA, Frank LG, Smale L. Patterns of Association among Female Spotted Hyenas (Crocuta crocuta). J Mammal. 1997 Feb 21;78(1):55–64.

46. Hofer H, Benhaiem S, Golla W, East ML. Trade-offs in lactation and milk intake by competing siblings in a fluctuating environment. Behav Ecol. 2016;27(5):1567–78.

47. Drea CM, Frank LG. 5. The Social Complexity of Spotted Hyenas. In: De Waal FBM, Tyack PL, editors. Animal Social Complexity [Internet]. Harvard University Press; 2003 [cited 2025 Mar 24]. p. 121–48. Available from: https://www.degruyter.com/document/doi/10.4159/harvard.9780674419131.c10/html

48. Gicquel M, East ML, Hofer H, Benhaiem S. Early-life adversity predicts performance and fitness in a wild social carnivore. J Anim Ecol. 2022 Oct;91(10):2074–86.

49. Watts HE, Holekamp KE. Ecological Determinants of Survival and Reproduction in the Spotted Hyena. J Mammal. 2009 Apr 14;90(2):461–71.

50. Höner OP, Wachter B, Hofer H, Wilhelm K, Thierer D, Trillmich F, et al. The fitness of dispersing spotted hyaena sons is influenced by maternal social status. Nat Commun. 2010 Aug 24;1(1):60.

51. Turner JW, Robitaille AL, Bills PS, Holekamp KE. Early-life relationships matter: Social position during early life predicts fitness among female spotted hyenas. Farine D, editor. J Anim Ecol. 2021 Jan;90(1):183–96.

52. Engh AL, Nelson KG, Peebles R, Hernandez AD, Hubbard KK, Holekamp KE. Coprologic Survey of Parasites of Spotted Hyenas (Crocuta crocuta) in the Masai Mara National Reserve, Kenya. J Wildl Dis. 2003 Jan;39(1):224–7.

53. East ML, Kurze C, Wilhelm K, Benhaiem S, Hofer H. Factors influencing *Dipylidium* sp. infection in a free-ranging social carnivore, the spotted hyaena (*Crocuta crocuta*). Int J Parasitol Parasites Wildl. 2013 Dec 1;2:257–65.

54. East ML, Otto E, Helms J, Thierer D, Cable J, Hofer H. Does lactation lead to resource allocation trade-offs in the spotted hyaena? Behav Ecol Sociobiol. 2015 May;69(5):805–14.

55. Benhaiem S, Dehnhard M, Bonanni R, Hofer H, Goymann W, Eulenberger K, et al. Validation of an enzyme immunoassay for the measurement of faecal glucocorticoid metabolites in spotted hyenas (Crocuta crocuta). Gen Comp Endocrinol. 2012 Sept;178(2):265–71.

56. Hofer H, East ML. Behavioral processes and costs of co-existence in female spotted hyenas: a life history perspective. Evol Ecol. 2003 July;17(4):315–31.

57. Hofer H, East ML. The commuting system of Serengeti spotted hyaenas: how a predator copes with migratory prey. I. Social organization. Anim Behav. 1993 Sept;46(3):547–57.

58. East ML, Höner OP, Wachter B, Wilhelm K, Burke T, Hofer H. Maternal effects on offspring social status in spotted hyenas. Behav Ecol. 2009;20(3):478–83.

59. Golla W, Hofer H, East ML. Within-litter sibling aggression in spotted hyaenas: effect of maternal nursing, sex and age. Anim Behav. 1999 Oct;58(4):715–26.

60. Frank LG, Glickman SE, Licht P. Fatal Sibling Aggression, Precocial Development, and Androgens in Neonatal Spotted Hyenas. Science. 1991 May 3;252(5006):702–4.

61. East ML, Hofer H. Male spotted hyenas (Crocuta crocuta) queue for status in social groups dominated by females. Behav Ecol. 2001 Sept 1;12(5):558–68.

62. Goymann W, East ML, Wachter B, Höner OP, Möstl E, Van’t Holf TJ, et al. Social, state-dependent and environmental modulation of faecal corticosteroid levels in free-ranging female spotted hyenas. Proc R Soc Lond B Biol Sci. 2001 Dec 7;268(1484):2453–9.

63. R Core Team. R: A Language and Environment for Statistical Computing. [Internet]. R Foundation for Statistical Computing, Vienna, Austria.; 2025. Available from: https://www.R-project.org/

64. Posit team. RStudio: Integrated Development Environment for R. [Internet]. Posit Software, PBC, Boston, MA.; 2025. Available from: http://www.posit.co/

65. Lüdecke D, Ben-Shachar M, Patil I, Waggoner P, Makowski D. performance: An R Package for Assessment, Comparison and Testing of Statistical Models. J Open Source Softw. 2021 Apr 21;6(60):3139.

66. Therneau, T. M. A Package for Survival Analysis in R [Internet]. 2020. Available from: https://CRAN.R-project.org/package=survival

67. Kassambara, A., Kosinski, M., Biecek, P.7. survminer: Drawing Survival Curves using ‘ggplot2’ [Internet]. 2024. Available from: https://rpkgs.datanovia.com/survminer/index.html

68. Hartig, F. DHARMa: residual diagnostics for hierarchical (multi-level/mixed) regression models [Internet]. 2016. Available from: https://cran.r-project.org/package=DHARM

69. Lüdecke D. ggeffects: Tidy Data Frames of Marginal Effects from Regression Models. J Open Source Softw. 2018 June 29;3(26):772.

70. Wickham H. ggplot2: Elegant Graphics for Data Analysis. Springer-Verl N Y [Internet]. 2016; Available from: https://ggplot2.tidyverse.org

71. Periago MV, Bethony JM. Hookworm virulence factors: making the most of the host. Microbes Infect. 2012 Dec;14(15):1451–64.

72. Seguel M, Gottdenker N. The diversity and impact of hookworm infections in wildlife. Int J Parasitol Parasites Wildl. 2017 Dec;6(3):177–94.

73. DeLong RL, Orr AJ, Jenkinson RS, Lyons ET. Treatment of northern fur seal (*Callorhinus ursinus*) pups with ivermectin reduces hookworm-induced mortality. Mar Mammal Sci. 2009 Oct;25(4):944–8.

74. Marcus AD, Higgins DP, Gray R. Epidemiology of hookworm (Uncinaria sanguinis) infection in free-ranging Australian sea lion (Neophoca cinerea) pups. Parasitol Res. 2014 Sept;113(9):3341–53.

75. Seguel M, Muñoz F, Navarrete MJ, Paredes E, Howerth E, Gottdenker N. Hookworm Infection in South American Fur Seal (*Arctocephalus australis*) Pups: Pathology and Factors Associated With Host Tissue Damage and Mortality. Vet Pathol. 2017 Mar;54(2):288–97.

76. Shoop WL. Vertical transmission of helminths: Hypobiosisand amphiparatenesis. Parasitol Today. 1991 Jan;7(2):51–4.

77. Traversa D. Pet roundworms and hookworms: A continuing need for global worming. Parasit Vectors. 2012 Dec;5(1):91.

78. Drake ED, Ravindran S, Bal X, Pemberton JM, Pilkington JG, Nussey DH, et al. Sex-specific effects of early-life adversity on adult fitness in a wild mammal. Proc R Soc B Biol Sci. 2025 Mar;292(2043):20250192.

79. Behnke JM. Structure in parasite component communities in wild rodents: predictability, stability, associations and interactions …. or pure randomness? Parasitology. 2008 June;135(7):751–66.

80. Graham AL. Ecological rules governing helminth–microparasite coinfection. Proc Natl Acad Sci. 2008 Jan 15;105(2):566–70.

81. Telfer S, Birtles R, Bennett M, Lambin X, Paterson S, Begon M. Parasite interactions in natural populations: insights from longitudinal data. Parasitology. 2008 June;135(7):767–81.

82. Woolhouse MEJ. Patterns in Parasite Epidemiology: The Peak Shift. Parasitol Today. 1998 Oct;14(10):428–34.

83. Cattadori IM, Boag B, Bjørnstad ON, Cornell SJ, Hudson PJ. Peak shift and epidemiology in a seasonal host–nematode system. Proc R Soc B Biol Sci. 2005 June 7;272(1568):1163–9.

84. Macpherson AJ, Geuking MB, McCoy KD. Immunoglobulin A: a bridge between innate and adaptive immunity. Curr Opin Gastroenterol. 2011 Nov;27(6):529–33.

85. Macpherson AJ, Geuking MB, Slack E, Hapfelmeier S, McCoy KD. The habitat, double life, citizenship, and forgetfulness of IgA. Immunol Rev. 2012 Jan;245(1):132–46.

86. Butler JE. Immunoglobulin diversity, B-cell and antibody repertoire development inlarge farm animals: -EN- -FR- -ES-. Rev Sci Tech OIE. 1998 Apr 1;17(1):43–70.

87. Veloso Soares SP, Jarquín-Díaz VH, Veiga MM, Karl S, Czirják GÁ, Weyrich A, et al. Mucosal immune responses and intestinal microbiome associations in wild spotted hyenas (Crocuta crocuta). Commun Biol. 2025 June 13;8(1):924.

88. Benhaiem S, Hofer H, Dehnhard M, Helms J, East ML. Sibling competition and hunger increase allostatic load in spotted hyaenas. Biol Lett. 2013 June 23;9(3):20130040.

89. Bonier F, Martin PR, Moore IT, Wingfield JC. Do baseline glucocorticoids predict fitness? Trends Ecol Evol. 2009 Nov;24(11):634–42.

90. Busch DS, Hayward LS. Stress in a conservation context: A discussion of glucocorticoid actions and how levels change with conservation-relevant variables. Biol Conserv. 2009 Dec;142(12):2844–53.

91. Dantzer B, Westrick SE, Van Kesteren F. Relationships between Endocrine Traits and Life Histories in Wild Animals: Insights, Problems, and Potential Pitfalls. Integr Comp Biol. 2016 Aug;56(2):185–97.

92. Defolie C, Merkling T, Fichtel C. Patterns and variation in the mammal parasite–glucocorticoid relationship. Biol Rev. 2020 Feb;95(1):74–93.

93. Romeo C, Wauters LA, Santicchia F, Dantzer B, Palme R, Martinoli A, et al. Complex relationships between physiological stress and endoparasite infections in natural populations. Ferkin M, editor. Curr Zool. 2020 Oct 1;66(5):449–57.

94. Friant S, Ziegler TE, Goldberg TL. Changes in physiological stress and behaviour in semi-free-ranging red-capped mangabeys (*Cercocebus torquatus*) following antiparasitic treatment. Proc R Soc B Biol Sci. 2016 July 27;283(1835):20161201.

95. Müller-Klein N, Heistermann M, Strube C, Morbach ZM, Lilie N, Franz M, et al. Physiological and social consequences of gastrointestinal nematode infection in a nonhuman primate. Behav Ecol. 2019 Apr 5;30(2):322–35.

96. Behringer V, Müller-Klein N, Strube C, Schülke O, Heistermann M, Ostner J. Responsiveness of fecal immunoglobulin A to HPA-axis activation limits its use for mucosal immunity assessment. Am J Primatol. 2021 Dec;83(12):e23329.

97. Cizauskas CA, Turner WC, Pitts N, Getz WM. Seasonal Patterns of Hormones, Macroparasites, and Microparasites in Wild African Ungulates: The Interplay among Stress, Reproduction, and Disease. Sreevatsan S, editor. PLOS ONE. 2015 Apr 15;10(4):e0120800.

98. Behie AM, Pavelka MSM. Interacting Roles of Diet, Cortisol Levels, and Parasites in Determining Population Density of Belizean Howler Monkeys in a Hurricane Damaged Forest Fragment [Internet]. Unpublished; 2014 [cited 2025 Jan 18]. Available from: http://rgdoi.net/10.13140/2.1.3426.3685

99. Hernandez SE, Strona ALS, Leiner NO, Suzán G, Romano MC. Seasonal changes of faecal cortisol metabolite levels in Gracilinanus agilis (Didelphimorphia: Didelphidae) and its association to life histories variables and parasite loads. Conserv Physiol [Internet]. 2018 Jan 1 [cited 2025 Jan 18];6(1). Available from: https://academic.oup.com/conphys/article/doi/10.1093/conphys/coy021/5055533

100. Touma C, Palme R. Measuring Fecal Glucocorticoid Metabolites in Mammals and Birds: The Importance of Validation. Ann N Y Acad Sci. 2005 June;1046(1):54–74.

101. Seeley KE, Proudfoot KL, Edes AN. The application of allostasis and allostatic load in animal species: A scoping review. Giroud S, editor. PLOS ONE. 2022 Aug 30;17(8):e0273838.

102. Martin LB. Stress and immunity in wild vertebrates: Timing is everything. Gen Comp Endocrinol. 2009 Sept;163(1–2):70–6.

103. Edwards PD, Lavergne SG, McCaw LK, Wijenayake S, Boonstra R, McGowan PO, et al. Maternal effects in mammals: Broadening our understanding of offspring programming. Front Neuroendocrinol. 2021 July;62:100924.

104. Mejía M, Gonzalez-Iglesias A, Díaz-Torga GS, Villafañe P, Formía N, Libertun C, et al. Effects of continuous ivermectin treatment from birth to puberty on growth and reproduction in dairy heifers. J Anim Sci. 1999;77(6):1329.

105. Blackwell AD, Tamayo MA, Beheim B, Trumble BC, Stieglitz J, Hooper PL, et al. Helminth infection, fecundity, and age of first pregnancy in women. Science. 2015 Nov 20;350(6263):970–2.

106. Wikenros C, Gicquel M, Zimmermann B, Flagstad Ø, Åkesson M. Age at first reproduction in wolves: different patterns of density dependence for females and males. Proc R Soc B Biol Sci. 2021 Apr 14;288(1948):rspb.2021.0207, 20210207.

107. Kutzer MAM, Armitage SAO. Maximising fitness in the face of parasites: a review of host tolerance. Zoology. 2016 Aug;119(4):281–9.

108. Neuhaus P, Broussard DR, Murie JO, Dobson FS. Age of primiparity and implications of early reproduction on life history in female Columbian ground squirrels. J Anim Ecol. 2004 Jan;73(1):36–43.

109. Nilsen EB, Brøseth H, Odden J, Linnell JDC. The cost of maturing early in a solitary carnivore. Oecologia. 2010 Dec;164(4):943–8.

110. Dickinson ER, Orsel K, Cuyler C, Kutz SJ. Life history matters: Differential effects of abomasal parasites on caribou fitness. Int J Parasitol. 2023 Apr;53(4):221–31.

111. Pigeon G, Pelletier F. Direct and indirect effects of early-life environment on lifetime fitness of bighorn ewes. Proc R Soc B Biol Sci. 2018 Jan 10;285(1870):20171935.

112. Pedersen AB, Fenton A. The role of antiparasite treatment experiments in assessing the impact of parasites on wildlife. Trends Parasitol. 2015 May;31(5):200–11.

113. Rózsa L, Reiczigel J, Majoros G. QUANTIFYING PARASITES IN SAMPLES OF HOSTS. J Parasitol. 2000 Apr;86(2):228–32.

